# On wing pattern and wing shape evolution in Giant Silk Moths: Lessons from wing development in Luna and Polyphemus Moths

**DOI:** 10.1101/2025.06.01.657247

**Authors:** Andrei Sourakov

**Affiliations:** McGuire Center for Lepidoptera and Biodiversity, Florida Museum of Natural History, University of Florida, Gainesville, FL, USA

**Keywords:** evo-devo, genetics, genotype, regulation, phenotype, *Wnt*, *wingless*, CRE, cis-regulatory elements, plasticity

## Abstract

Heparin and dextran sulfate were used to influence wing development in the Luna Moth and the Polyphemus Moth. The experiments led to alterations not only of wing patterns, but also of wing shapes in both species, albeit in different ways. Dextran sulfate led to contractions of the wing pattern elements that were expanded by heparin in the Polyphemus Moth. Transformations of the dorsal wing pattern of the Luna Moth under the influence of heparin were dramatic and demonstrated the hidden potential for creating color patterns by contracting and expanding wing pattern elements. For the first time, polysaccharides have been shown to have an effect on wing shape: heparin caused significant expansion of the hindwing tails in the Luna Moth, while dextran sulfate caused a narrowing of the wings in the Polyphemus Moth. This study aims to provide insight into homologies in wing pattern elements, such as eyespots, across the Lepidoptera order, in general, and Saturniidae, in particular, and contributes to understanding phenotypic diversity and plasticity within the genera *Actias* and *Antheraea*.

---

Butterfly wing patterns have received a lot of attention from the developmental biology community in the last 20 years (e.g., Marcus, 2021). For example, hundreds of evo-devo studies by a variety of research groups have been enabled by laboratory cultures of *Bicyclus anynana* satyrine butterflies maintained across many institutions. These populations enabled researchers to experimentally probe genetic controls of prominent eyespot characteristics, like size, location, and color, and to better understand their function within butterfly morphology (e.g., Beldade and Monteiro, 2021).

Eyespots have originated numerous times independently across Lepidoptera (e.g., Hanotte et al., 2024) and across other groups of insects, such as Coleoptera or Neuroptera (e.g., Machado et al., 2021). While the majority of eyespot studies have been conducted on the nymphalid family of butterflies, moth eyespots have also been examined, albeit more rarely, leaving numerous questions unanswered. Saturniidae, or the Giant Silk Moths, are an ideal group for experimentation, as they are large, well-understood taxonomically, diverse morphologically and evolutionarily, and are relatively easy to rear. In a recent study on the influence of heparin on the eyespots of the Io Moth (*Automeris io*) and the Polyphemus Moth (*Antheraea polyphemus*), it was demonstrated that concentrically organized circles around the M_2_-M_3_ cross vein are in fact comprised of non-homologous elements, despite their similarity in appearance and their shared function of warding off predators (Sourakov and Shirai, 2020).

Heparin, and dextran sulfate as its antagonist, can upregulate and downregulate, respectively, *Wnt* signaling, as was first demonstrated by Serfas and Carroll (2005) and used numerous times since (e.g., Martin and Reed, 2014; Mazo-Vargas, 2020). As a result, injecting these substances into developing immature stages during crucial stages of wing pattern formation (late prepupa to early pupa) can shed light onto the mysteries of specific wing patterns and their homologies (e.g., Sourakov et al., 2022; Mazo-Vargas et al., 2024). Recent findings from experiments by Otaki and Nakazato (2022) hint at the mechanism, demonstrating that inducers of wing color pattern modification may act on the chitin of pupa and thus influence morphogenic signals that determine color patterns.

Experiments with wing pattern modifiers can potentially offer evo-devo researchers help in planning follow-up experiments aimed at determining gene expression in the forming wing (Anyi Mazo-Vargas, pers. com., 2025). For example, such experiments by Sourakov and Shirai (2020) with *Antheraea polyphemus* were followed up by RNA-seq expression experiments on this species, comparing wild-type and heparin-treated individuals (Pomerantz, 2021, pp. 97-100).

Here, I report new results of experiments involving wing pattern modifiers conducted on the Luna Moth (*Actias luna*) and the Polyphemus Moth (*Antheraea polyphemus*). Surprisingly, it was found that not only wing pattern changes but also wing shape changes can be induced by using heparin and dextran sulfate injections. Thanks to predation experiments, we now have great insight into the function of hindwing tails in the Luna Moth (e. g., Barber et al., 2015), so it is my belief that wing shape evo-devo research will prove especially interesting when conducted on saturniids. It is my hope that the results presented here will assist others in the Lepidoptera research community in looking for genes responsible for wing shape modification in this moth family.

## Materials and Methods

Experiments were conducted between 2020 and 2025, on several laboratory broods of *Actias luna* and *Antheraea polyphemus* that originated from females collected in Gainesville, Florida. Caterpillars were raised on Sweet Gum (*Liquidambar styraciflua*) and Water Oak (*Quercus nigra*), respectively. Over 500 *A. luna* moths were raised to adult stage, with many injected with solutions of heparin or dextran sulfate as late prepupa or early pupa. As controls, individuals from the same broods were either injected with water or Ringer solution or left unmanipulated. Additionally, individuals injected with lower doses of heparin and dextran sulfate served as “controls” for the individuals injected with higher concentrations. All adult specimens were mounted and received unique MGCL voucher numbers. They will be deposited into the collection of the McGuire Center for Lepidoptera (MGCL) and Biodiversity at the Florida Museum of Natural History. While *A. luna* served as the main experimental targets, two dozen *A. polyphemus* moths underwent similar experiments and serve as a contrast to the transformations imparted on *A. luna*.

The injections were conducted using methods described earlier by Sourakov (2020) and Sourakov & Shirai (2020) using sodium salt from porcine (MP Biomedicals, Inc.) dissolved in distilled water to concentrations ranging from 5 to 55% (by weight). Dextran sulfate sodium salt from *Leuconostoc* spp. (Sigma, Inc.) was dissolved in distilled water or Ringer solution to concentrations ranging from 7 to 31% (by weight). Injections were conducted at room temperature at different quantities and at different intervals before and after pupation using 0.3-ml hypodermic syringe. Injections were conducted subcutaneously in prepupae or in the upper abdomen dorsally (mostly) or wing compartments (occasionally) in pupae.

## Results and Discussion

### Luna Moth: wing shape and pattern changes

Experiments with heparin injections were conducted to test the eyespots of the Luna Moth in the same way in which it was previously done with the Io Moth and the Polyphemus Moth by Sourakov and Shirai (2020). Unexpectedly, these manipulations led to alterations of not only the wing pattern but also of the wing shape of the Luna Moth. For instance, the wing tails became as much as 1.8 times wider in some of the transformed individuals compared to control individuals (Fig. 1). Dextran sulfate injections, on the other hand, contracted the wings, causing deformities.

**Fig. 1.**
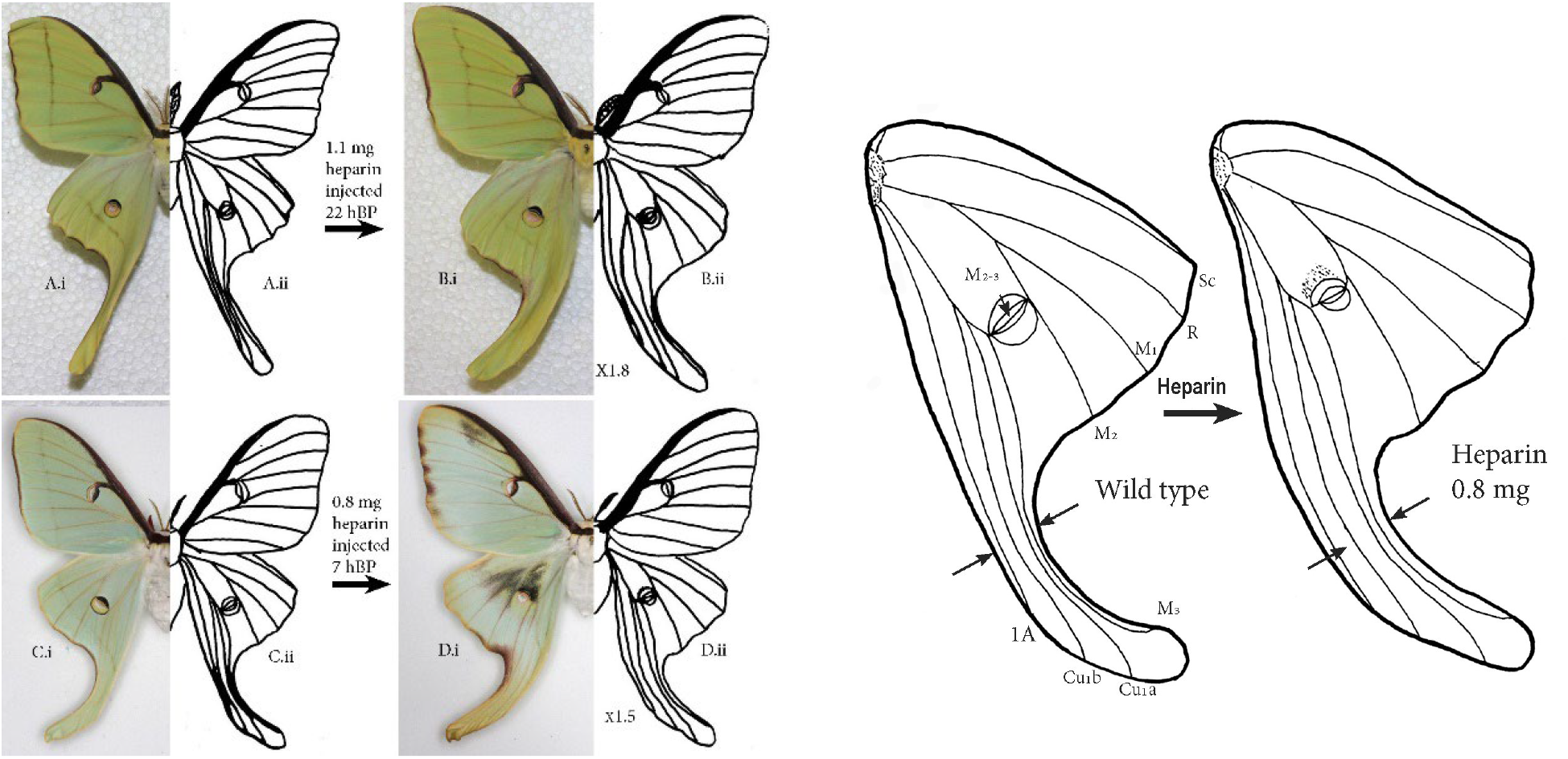
Luna Moth transformation under the influence of heparin. (**Left**) Heparin caused 1.5-1.8 times expansion of the width of the hindwing tails, depending on the timing of the injection as well as the heparin quantity: (**A**) wild-type male and (**B**) its sibling male injected with heparin 22 hours before pupation; (**C**) wild-type female and (**D**) its sibling female injected with heparin 7 hours before pupation. (**Right**) Hindwings of individuals C and D showing wing venation. Sc - subcostal, R - radial, M - medial, Cu - cubital, A - anal veins; arrows indicate heparin-induced change in wing tail width.

Up to this point, the Marginal Band Symmetry system (MBS) as well as the Central Symmetry System (CSS) have been shown to be positively affected by heparin injections and negatively affected by dextran sulfate injections (e. g., Serfas and Carroll, 2005; Martin and Reed, 2014). While in the wild-type Luna Moth, most symmetry systems are not well-pronounced, the transformations induced by heparin allowed some of them to be revealed by greatly expanding them, creating much more patterned individuals (Figs. 1, 4). Expansion of the hindwing area and tails in individuals injected with high concentrations of heparin as prepupae/early pupae may suggest a possible broader role of *Wnt*-ligand genes in this species but may also have a completely different mechanism behind it.

### Polyphemus Moth: wing shape and pattern changes

While heparin-induced changes to the wing pattern were expected based on previous works on this species (Sourakov and Shirai, 2020, Pomeranz, 2021), the dextran sulfate results are new, and offer additional evidence that these two substances work as antagonists of each other if injected during wing pattern formation. From photos of nine siblings arranged side by side (Fig. 2 (left)), one can easily observe that they are different, with eyespot elements of heparin-injected individuals expanding, and of the dextran sulfate-injected individuals contracting. The hindwing eyespot width (measured in relation to the forewing length) shrunk significantly (P=0.0017) in dextran sulfate-injected individuals, with the borders of the eyespots formed by the outer black ring (indicated as “R” in Fig. 2) much more tightly delimited rather than “bleeding” into the surrounding ground color as in the wild-type individuals. This black ring greatly expands under the influence of heparin. The same observations apply to other eyespot elements (“G”, “B”, and “W” in Fig. 2 (right)). Changes were also imparted by heparin injections on the dorsal forewing pattern of the Polyphemus Moth (see Fig. 4), and they will be further discussed below.

**Fig. 2.**
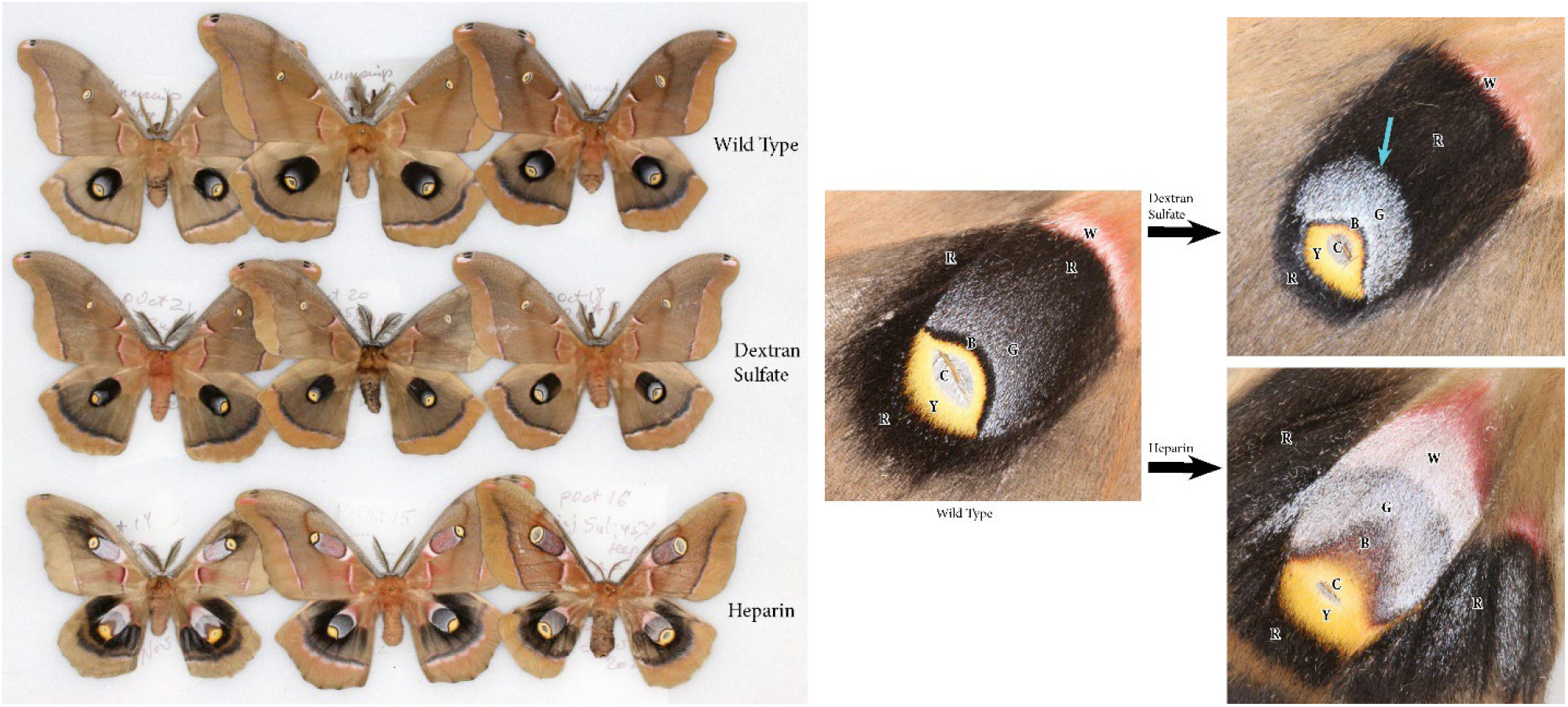
Polyphemus Moth transformation under the influence of heparin and dextran sulfate. (**Left**) Wild-type, dextran sulfate, and heparin-affected sibling individuals form three distinct phenotypic groups. (**Right**) Effect of heparin and dextran sulfate on the dorsal hindwing eyespots; Y, B, G, R, and W elements forming the eyespot expand under the influence of heparin (bottom right) and contract under the influence of dextran sulfate (top right) as compared to the wild type (left). It appears that the clear area “C” surrounding the M_2_-M_3_ cross vein in the eyespot center does not expand under the influence of heparin, indicating that it is formed by a scale-less wing membrane or flattened expanded cross vein, unlike the rest of the wing pattern, which is created by scales of different colors.

Less obvious than in the Luna Moth were the changes to the wing shapes in the Polyphemus Moth. However, measurements of forewing length and width among males (two subgroups for which there was a sufficient sample size) demonstrate that their ratio was greater in dextran sulfate-injected individuals (N=4) than in their unmanipulated siblings (N=7) (P=0.0164), indicating that the wing becomes narrower when subjected to dextran sulfate treatment. The same effect was observed in the hindwing (P=0.0359).

### Additional details

Achieving changes in the wing pattern of *Actias luna* required much higher heparin concentrations than in previous experiments with other species. The changes, consisting of a significant expansion in the dark scales of the hindwing eyespot and of the normally very narrow, burgundy-colored marginal band, were eventually achieved in a female shown in Fig. 1, which received two injections, 0.4 mg of heparin each, as a late prepupa around 7 hours before pupation (hBP). In addition to wing pattern changes, the width of hindwing tails was modified in this and another specimen that was injected as a prepupa with 1.1. mg of heparin at 22 hBP (Fig. 1, top left). That specimen showed only minor modifications of the wing pattern consisting of a “smudging” of the marginal band, perhaps indicating that the injection was made too early in the wing development for the rest of the wing pattern to be affected.

Lower heparin quantities administered to prepupae and pupae achieved no results, leading to the emergence of wild-type individuals. Although higher doses of heparin corresponded to an increased mortality rate, eventually a relatively high survival and transformation rate was achieved. In total, a noticeable wing pattern alteration was achieved in 86 individuals, some very dramatic (e. g., Fig. 4). In the Luna Moth, dextran sulfate causes contraction of wings by seemingly impeding hemolymph flow, especially via some of the veins, and it is important to note that this known heparin antagonist has a different effect on the Polyphemus Moth (see Figs. 2, 4).

### Evolutionary implications

As discussed in the introduction, hindwing tails in the genus *Actias* are very important for their survival and, as a result, are very pliable. The tail length is variable interspecifically, and this difference was suggested to depend on the predation environment by Rubin et al., (2025). There are also a few cases where there is significant sexual dimorphism, such as displayed by *A. isabellae*, whose females possess hindwing tails that are shorter and wider than their male counterparts (Fig. 3). It would be interesting if it turned out that the same genetic pathways that are responsible for wing shape differences in *A. isabellae* (and some other sexually dimorphic species of *Actias*) are also the ones that are responsive to heparin in the present experiments on the Luna Moth.

**Fig. 3.**
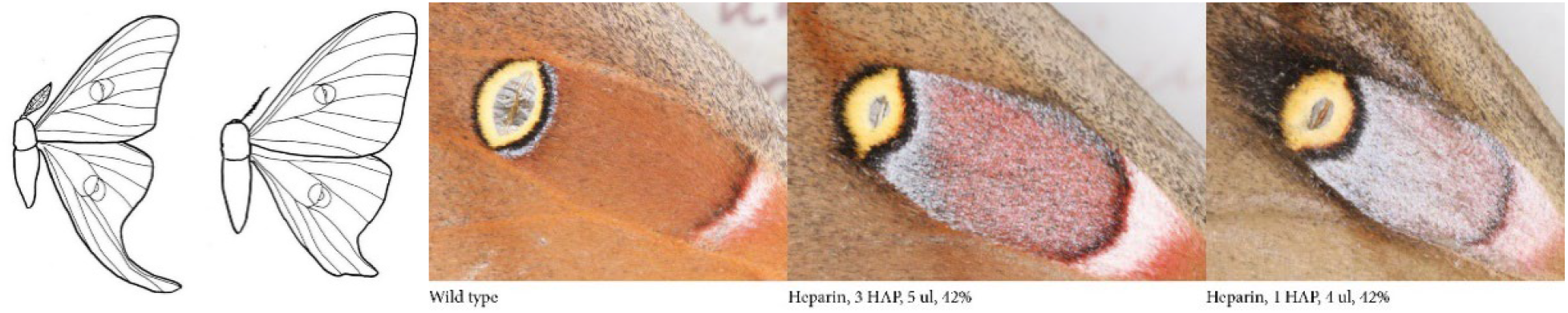
(**Left**) Wing venation of the Spanish Moon Moth, *Actias isabellae*. Male (on the left) and female demonstrate different hindwing shapes, indicating differences in selective pressure on this feature in different sexes. (**Right**) From cryptic to aposematic: The dorsal forewing eyespot of the Polyphemus Moth transformed under the influence of heparin.

The Polyphemus Moth normally rests with its wings closed, exposing the ventral wing surface. If disturbed, the moth opens its wings and brings forewings upward, exposing the hindwing eyespots to ward off a predator. So, the dorsal eyespot in this species normally simulates a hole in a dry leaf which the moth resembles when at rest. However, as shown in Fig. 3, this modest-looking eyespot still contains compressed aposematic pattern-forming potential. This experiment demonstrates that a simple upregulation of certain wing pattern genes may be sufficient to change the function of the eyespot from cryptic to aposematic.

With its dorsal wing pattern (as well as its shape), the Polyphemus Moth mimics dry foliage when at rest. The dorsal fore- and hindwing patterns were changed by heparin and dextran sulfate (Fig. 4). Based on the current knowledge of heparin action on a developing wing, combined with results of CRISPR experiments (Mazo-Vargas et al., 2022), the present experiments demonstrate how up- and down regulation of certain pathways can create individuals within a population (and even within a brood) that are phenotypically varied. In the case of the Polyphemus Moth, this means that avian predators would be unable to develop a search pattern for a particular “leaf” they may otherwise come to associate with a tasty meal.

**Fig. 4.**
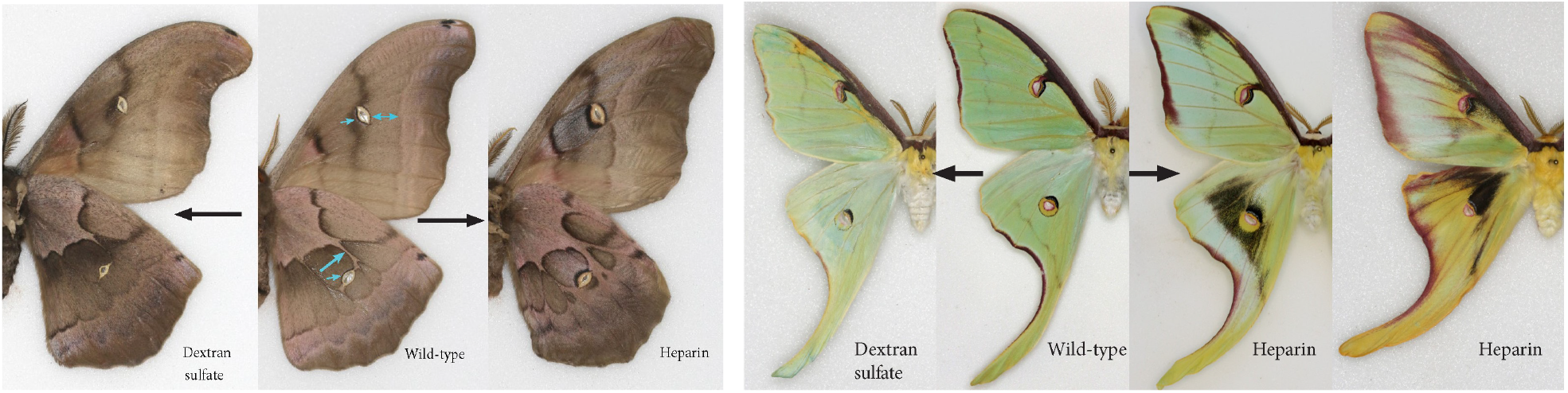
(**Left**) Ventral wing pattern transformations in the Polyphemus Moth resulting from heparin and dextran sulfate injections during wing pattern development in the early pupal stage. The wild-type (middle) is intermediate in its wing pattern between the dextran sulfate and heparin-affected individuals, as can be observed by focusing on the elements indicated with arrows. Dextran sulfate caused contractions of those elements which are expanded under the influence of heparin. (**Right**) Dorsal wing pattern transformations of Luna Moth males under the influence of heparin demonstrates the hidden potential for creating color patterns in related species by upregulating development of normally contracted wing pattern elements. In contrast with the Polyphemus Moth, dextran sulfate contracted wing veins in this species, negatively affecting wing expansion after emergence from the pupa.

Likewise, in the Luna Moth, from highly compressed, but nevertheless present wing pattern elements such as their normal modest eyespots and narrow, burgundy-colored wing margins, a completely novel patterns were created by upregulating some of the pathways. Heparin treatments have also revealed, especially in males (Fig. 4) the complexity of green color in this species, which seems to be a combination of the light base color and the overlaying green scales.

## Acknowledgments

Laboratory broods of Luna and Polyphemus moths used in these experiments were started from eggs shared with me by Kevin Burnette, Bert Foquet, Eric Andreson, and Matthew Standridge. I am especially indebted to Eric Anderson and Alexandra Shapiro for looking after caterpillars in my absence and to Keith Willmott for sharing many sweetgum branches from his yard over the years. Many thanks go to my daughter, Alexandra, for proofreading this manuscript and offering helpful suggestions.

